# Dual-color expansion microscopy of membrane proteins using bioorthogonal labeling

**DOI:** 10.1101/2025.10.10.680277

**Authors:** Steven Edwards, Birthe Meineke, Sebastian Bauer, Hans Blom, Simon Elsässer, Hjalmar Brismar

**Author notes:** Correspondence should be addressed to H.B.

## Abstract

Site-specific incorporation of non-canonical amino acids (ncAAs) combined with bioorthogonal click chemistry provides a powerful tool for fluorescent protein labeling, overcoming the linkage error inherent to antibody-based probes. In this study, we present the development of dual-color super-resolution imaging utilizing ncAA labeling together with expansion microscopy (ExM). After optimizing the labeling procedures and fluorophore selection, we visualize and resolve the nanoscale distribution of Na,K-ATPase α_1_ and β_1_ subunits in expanded HEK 293T cells. We validate our approach by super-resolution STED imaging of ncAA labeled β_1_ subunit in unexpanded cells. This work establishes a robust framework for multiplexed, high-resolution imaging and suggests that the combination of ncAA labeling with ExM has the potential to push biological imaging toward Ångström-level resolution.

## Introduction

Deciphering the nanoscale organization of membrane proteins is fundamental to understanding cell biology, yet this remains a significant technical challenge due to the inherent difficulty in achieving labeling without spatial distortions and access to super-resolution microscopy methods. A key protein of interest is the Na,K-ATPase (NKA), a ubiquitous integral membrane protein essential for maintaining ion gradients in virtually all eukaryotic cells (1). While NKA is known to be a heterodimer composed of α and β subunits, its higher-order spatial arrangement within the plasma membrane is not fully understood (2–6). Determining whether NKA functions as individual heterodimers or as part of larger oligomeric complexes requires imaging technologies that can resolve molecules at a scale well below the diffraction limit of light (7).

To address this challenge, super-resolution microscopy techniques, such as STED and SMLM, have become valuable tools. However, the ultimate resolution of these methods is dependent on the labeling strategy (8). Traditional antibody based immunolabeling can introduce a “linkage error” of tens of nanometers due to the physical size of the antibodies, obscuring the true location of the target protein. Self-labeling tags like SNAP and Halo have been introduced as attractive alternatives, but there are reports of inconsistent labeling efficiency using those approaches due to varying cellular environments (9).

A more precise alternative is the site-specific incorporation of non-canonical amino acids (ncAAs) via genetic code expansion (GCE) (10). This technique allows for the placement of a small chemical handle at a specific site within a protein. A subsequent bioorthogonal “click chemistry” reaction can then attach an organic fluorophore with minimal linkage error, enabling a more precise representation of the protein’s location (11).

In parallel, expansion microscopy (ExM) has emerged as an attractive and accessible super-resolution technique (12). By physically enlarging the biological specimen within a swellable hydrogel, ExM makes it possible to achieve nanoscale resolution using conventional, diffraction-limited microscopes (13,14).

In this study, we combine these technologies, presenting a workflow that integrates GCE-based bioorthogonal labeling with ExM for dual-color super-resolution imaging. By targeting the α_1_ and β_1_ subunits of NKA, we visualize the enzyme in expanded HEK 293T cells, providing new insights into its nanoscale distribution (7,15,16). We further validate our approach through a qualitative comparison with STED microscopy. This work establishes a robust framework for multiplexed, high-resolution imaging and demonstrates how the synergy between ncAA labeling and ExM can advance biological nanoscale imaging.

## Methods

### Cell culture

HEK293T cells were passaged every 3-4 days and cultured at 37°C, 80% humidity and 5% CO2 in Dulbecco’s modified Eagle’s medium (DMEM, Sigma-Aldrich) media supplemented with 10% FBS and 1% penicillin-streptomycin. For transfection, cells were seeded at a density of 25 000 cells/cm^2^ onto 10 mm diameter #1.5 coverslips pre-coated with Poly-L-Lysine (P4707; Sigma-Aldrich). Cells were allowed to grow for 24 hours prior to transient transfection.

### DNA constructs and transfection

Plasmids for EF1 promoter-controlled expression of the NKA α_1_ and β_1_ subunits (rat) containing amber (TAG) or ochre (TAA) stop codons were generated based on pAS (#140008 Addgene). Point mutation for α_1_ T121TAG, β_1_ L64TAA and β_1_ L64TAG were introduced by two step PCR from the wild type sequences (3). To facilitate dual stop codon suppression the four tandem h7SK-PylT repeats were exchanged for a total of eight tRNA genes: four tandem repeats coding for *Mx1201/G1* hybrid tRNA^Pyl^ _CUA_ A41AA C55A (hyb*) controlled by h7SK promoter, and four tandem repeats coding for U6 promoter controlled *Mma* tRNA^Pyl^ variant M15 with a UUA anticodon (M15_UUA_). The two aminoacyl-tRNA synthetases (aaRS) used were pAS_4xhybPylT A41AA C55A FLAG-G1 PylRS Y125A (#154773; Addgene) for amber suppression and pAS_4xU6-PylT M15 (UUA) FLAG-Mma PylRS (#154774; Addgene) for ochre suppression (17).

Cells were transiently transfected with 500 ng DNA/cm^2^ with a 9:1 NKA α/β 1:aaRS ratio. TransIT-LT1 (MIR2305; Mirus) transfection reagent was used with Opti-MEM I reduced serum medium (31985062; Thermo Fisher Scientific) according to manufacturer’s instructions. In single labeling experiments, a WT NKA α_1_ or β_1_ subunit was co-transfected (1:1) with the mutated subunit.

The ncAAs axial trans-cyclooct-2-ene–lysine (TCO*K, #SC-8008; SiChem) and N-Propargyl-L-lysine (ProK, #HAA2090; Iris Biotech) were added to the cell culture media at the same time as transfection, the cells were then grown for 48 h to allow sufficient protein expression prior to labeling and fixation. Unless stated otherwise, final concentrations were 0.25 mM ProK and 0.1 mM and TCO*K, respectively.

### Click Chemistry

ProK was labeled using Cu(I)-catalyzed azide-alkyne cycloaddition (CuAAC) reaction using 1 µM dye AF647/488-picolyl azide (Jena Bioscience) in a buffer containing 50 µM CuSO_4_ (Copper(II) sulfate pentahydrate, #209198; Sigma-Aldrich), 250µM THPTA (Tris(3-hydroxypropyltriazolylmethyl)amine, #762342; Sigma-Aldrich) and 2.5 mM L-ascorbic acid (#A5960; Sigma-Aldrich) in PBS (18). CuSO_4_ and THPTA were vortexed together and kept on ice for 5 minutes before adding freshly dissolved L-Ascorbic acid, dye, and incubated for 10 minutes on ice. The reaction mixture was added to cells, and the reaction was allowed to proceed at room temperature for 5 minutes on live cells, 30-60 minutes on fixed cells, and 3hours on expanded gels.

TCO*K was labeled by using strain-promoted inverse electron-demand Diels-Alder cycloaddition (SPIEDAC) with 1 µM AF488/647-tetrazine (Click Chemistry Tools) or Abberior STAR 635/STAR RED-tetrazine (Abberior GmbH). For live cells, the dye was added directly to the PBS, and the cells were incubated for 30 minutes at 37°C. For fixed cells, the dye was mixed with PBS, and the cells were incubated at RT for 1 hour. For expanded gels, the dye was mixed in PBS, and the gels were allowed to incubate for 3 hours at room temperature.

### Expansion

Cells were fixed in 4% paraformaldehyde (PFA) for 15 minutes at RT prior to incubation in a humidified chamber at 37°C in AA/FA solution (30% Acrylamide (AA) and 4% Formaldehyde (FA)). Cells were washed 3×5min in PBS. 35 µL of monomer solution (7% sodium acrylate (SA), 20% AA, 0.05% N,N’-methylen-bisacrylamide (Bis-AA), 0.5% Ammonium persulfate (APS), and 0.5% N,N,N′,N′-Tetramethylethylenediamine (TEMED) in PBS was placed on glass slide covered with parafilm. The slide was placed on ice to create a cooled hydrophobic gelation surface. The coverslip was placed upside down onto the monomer solution and left for 5 minutes on ice before being moved to a humidified chamber at 37°C for 1.5 hours to polymerize. This coverslip and gel were removed from the parafilm and dropped into a denaturation buffer (200nM SDS, 200 mM NaCl, and 50 mM TRIS, pH 9.0) at 95°C for 1hour. The gels were allowed to fully expand by washing them several times in deionized water. The diameter of the fully expanded gel was measured and divided by the coverslip diameter (10 mm) to calculate the expansion factor.

### Mounting

Fully expanded gels were mounted on a glass-bottom dish (P35G-1.5-20-C; MatTek Corporation) precoated with Poly-L-Lysine (P4707; Sigma-Aldrich). Gels were covered with 1.5% UtraPure low melting point agarose (Invitrogen) in deionized H_2_O to support the gel during imaging. After polymerisation of the agarose, the dish was filled with deionized H_2_O.

### Imaging

Images of expanded gels were acquired on a Zeiss LSM 980 with Airyscan detection using a 40 × 1.2 NA water immersion objective. Images of fixed cells (Figure 1) were acquired using a Zeiss LSM 980 confocal with a 63 × 1.4 NA oil immersion objective. Confocal and STED images of fixed cells (Figure 2) were acquired using a using a Leica SP8 3X microscope with a 100 × 1.4 NA oil immersion objective.

**Figure 1.**
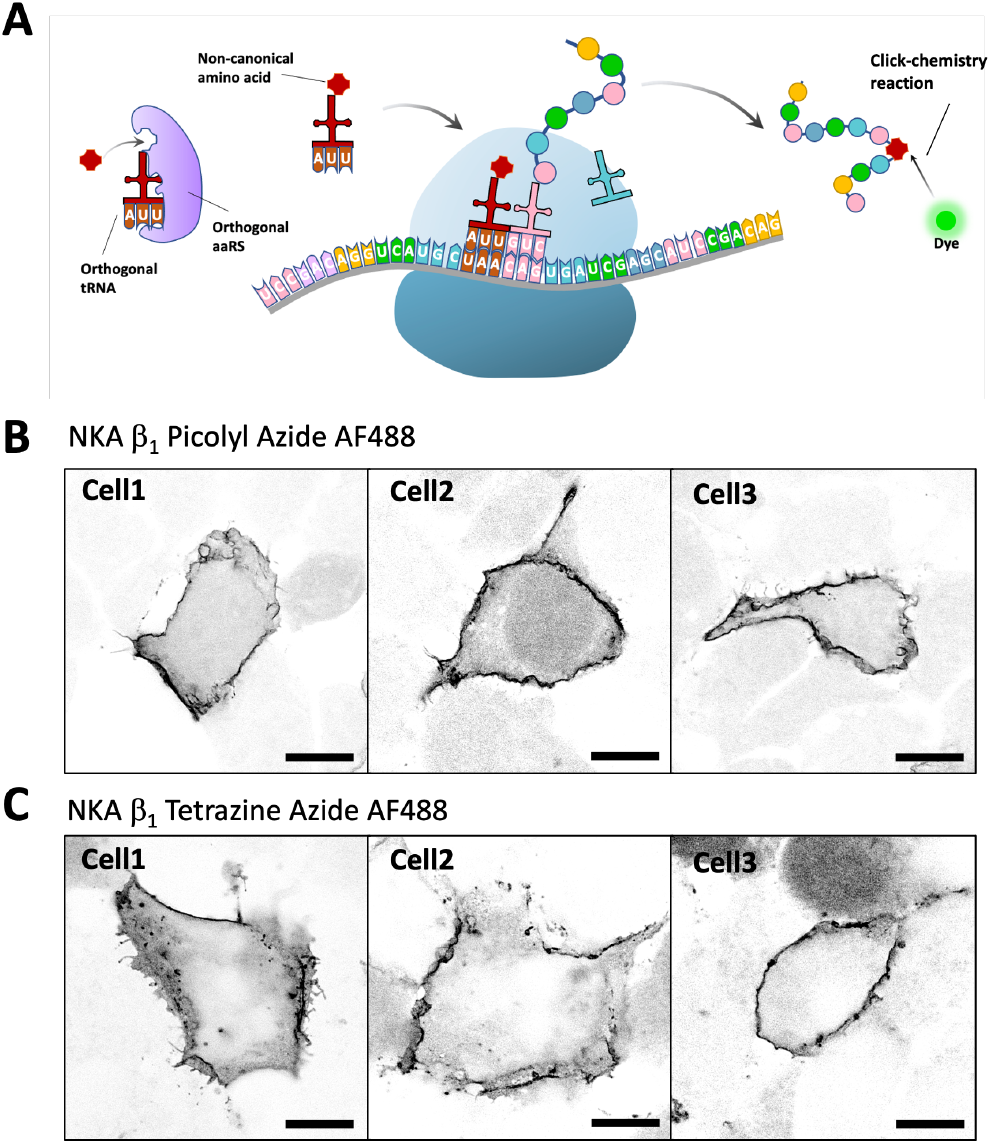
Incorporation of ncAA by genetic code expansion and their selective labeling by CuAAC and SPIEDAC click reactions. A schematic of genetic code expansion using an orthogonal tRNA/aaRS pair for site specific incorporation of ncAAs into proteins and fluorescent labeling using click chemistry (A). Confocal image of NKA β1 L64ProK CuAAC labeled with AF488 picolyl azide in the plasma membrane of fixed HEK239T cells (B). SPIEDAC labeling of NKA β1 L64TCO*K subunit in the plasma membrane of fixed HEK239T cells (C). Scale bars, 10 μm.

**Figure 2.**
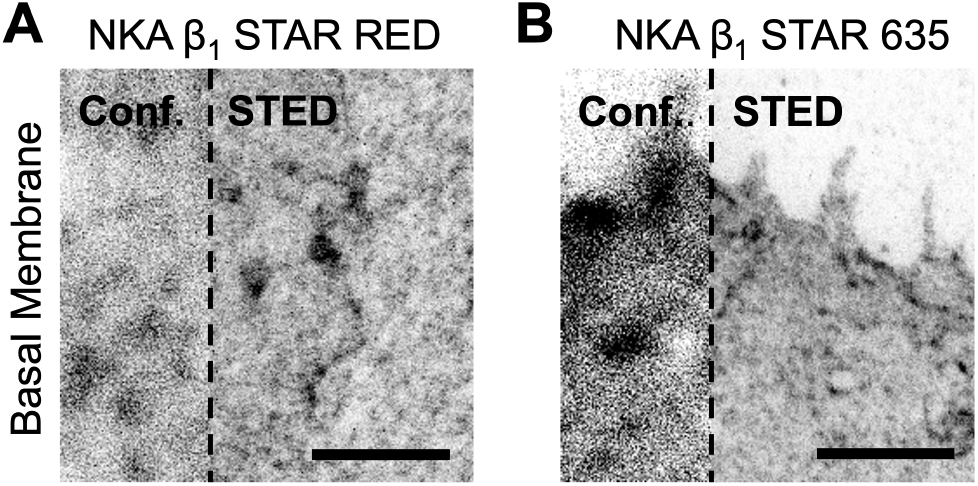
Super resolution STED microscopy of NKA β_1_ in the plasma membrane. SPIEDAC labeling of the NKA β1 L64TCO*K on the surface of HEK239T cells, fixed and imaged using confocal and STED microscopy. Abberior STAR RED tetrazine (A) and Abberior STAR 635 tetrazine (B) both showed labeling in the basal membrane. Scale bar 2 μm.

### Image analysis

To determine the size of individual NKA clusters, we identified local intensity maxima within the fluorescence images. The two-dimensional (2D) intensity profile surrounding each maximum was then fitted to a 2D Gaussian function. We defined the cluster diameter as the Full Width at Half Maximum (FWHM) of the resulting Gaussian fit.

## Results

### Genetic code expansion and click labeling of the NKA subunit β_**1**_ **in the plasma membrane**

Genetic code expansion via stop codon suppression allows the expression of ncAA containing proteins. Introduction of an orthogonal tRNA/aminoacyl-tRNA synthetase pair and its cognate ncAA recodes the stop codon and can lead to incorporation of the ncAA in that position, instead of translation termination (Figure 1A). We used genetic code expansion with amber (TAG), or ochre (TAA) stop codon suppression to introduce ncAAs with clickable functional groups into the extracellular region of the NKA β_1_ and α_1_ subunits. For ochre codon suppression we used *M. mazei (Mma)* pyrrolysyl-tRNA (tRNA^Pyl^) variant M15_UUA_/pyrrolysyl-tRNA synthetase (PylRS) pair which efficiently incorporates N-propargyl-L-lysine (ProK) (13). ProK can be specifically labeled with a picolyl azide modified dye using Cu(I)-catalyzed azide-alkyne cycloaddition (CuAAC).

To demonstrate successful incorporation of the ncAA and transport of NKA to the plasma membrane, HEK293T cells were co-transfected with NKA β_1_ L64TAA mutant, the M15_UUA_/*Mma*PylRS pair, and an NKA α_1_ WT to improve membrane insertion of the protein. The cell culture media was supplemented with ProK and cells were allowed to grow for 48 hours before click-labeling with AF488-picolyl azide, fixation and imaging using confocal microscopy. Fluorescent signal was detected in the plasma membrane of the cells (figure 1B), indicating expression, membrane insertion and click labeling of NKA β_1_ L64ProK.

We designed and produced a NKA β_1_ L64TAG mutant which was transfected with its amber suppressor hyb*/*G1*PylRS*YA* pair for TCO*K incorporation. TCO*K can be labeled by SPIEDAC click-chemistry with tetrazine modified dyes. The cell culture media was supplemented with TCO*K and cells were allowed to grow 48 hours before labeling with AF488-tetrazine, fixation and imaging using confocal microscopy. Once again, fluorescent signal was detected in the plasma membrane of the cells (Figure 1C). The AF488 dye used for click-labeling is membrane impermeable and should therefore only label the ncAA that are located extracellularly. However, when using SPIEDAC click-labeling, we occasionally observed unspecific background staining in cells which did not have a specific plasma membrane labeling (figure 1C).

### Super resolution STED microscopy of NKA β_1_ in the plasma membrane

To investigate the nanoscale distribution of the NKA β_1_ subunit, we first used STED super-resolution microscopy. Given that STED imaging requires bright and highly photostable fluorophores, we labeled the β_1_ subunit with either Abberior STAR 635-tetrazine or Abberior STAR RED-tetrazine dyes. As both dyes are membrane-impermeable, labeling was successfully restricted to β_1_ subunits located in the plasma membrane of the cells (Figure 2A, B). This is the first demonstration of successful use of STED optimized dyes for click-labeling.

The resulting STED images showed a non-homogenous distribution of NKA β_1_ at the cell membrane, which is consistent with previous reports (7).

### Expansion microscopy enables super resolution imaging of NKA β_1_ in the plasma membrane on diffraction limited microscopes

As an alternative approach to achieve super-resolution imaging, we utilized expansion microscopy (ExM), a technique that allows for nanoscale imaging on diffraction-limited microscopes. For this method, HEK293T cells were co-transfected to express NKA β_1_ L64ProK, which was subsequently click-labeled with AF488-picolyl azide. Proteins in the sample were then cross-linked into a swellable hydrogel. Following denaturation in an SDS containing buffer, the hydrogel was expanded in deionized water, resulting in an isotropic physical expansion of approximately 4.3-fold (Figure 3A).

**Figure 3.**
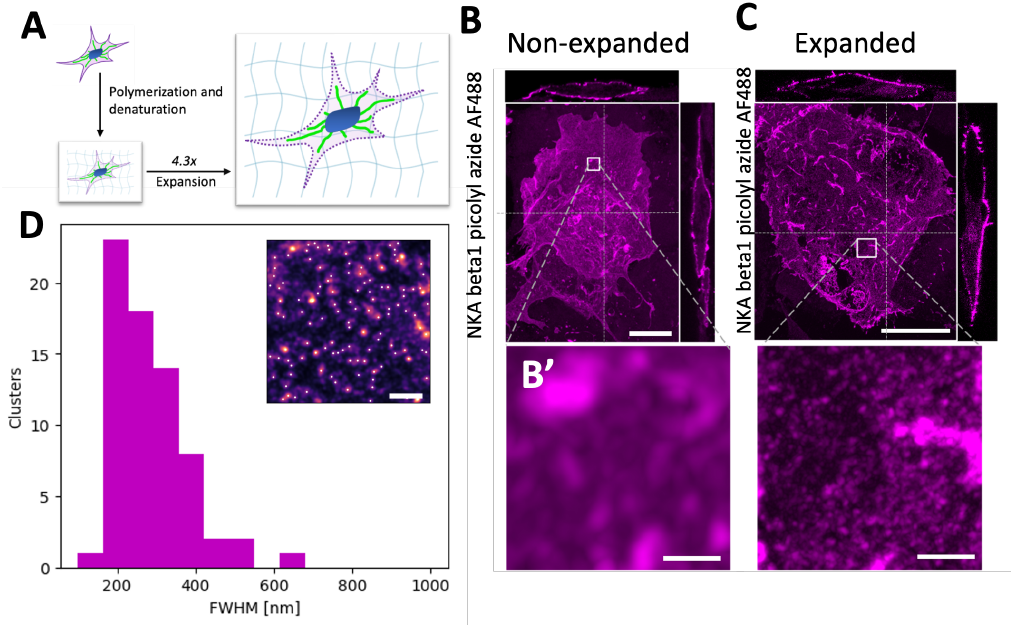
Expansion microscopy enables super resolution imaging of NKA β_1_ in the plasma membrane on diffraction limited microscopes. Schematic representation of the expansion microscopy process, a fixed cell is embedded into a swellable polyelectrolyte gel and denatured at 95 °C in an SDS containing buffer. The gel is immersed in deionized water to fully expand approximately 4.3-fold (A). Confocal z-stack (MIP) of non-expanded HEK293T cells expressing NKA β1 L64ProK labeled with AF488-picolyl azide by CuAAC. Orthogonal views are shown for the positions marked with a dashed line (B). Confocal z-stack (MIP) of expanded HEK293T cells expressing NKA β1 L64ProK labeled AF488-picolyl azide by CuAAC (C). Distribution of cluster sizes in the expanded cells. Clusters are identified by a spot analysis finding local intensity maxima and fitting to 2D Gaussian. Inset is a magnified view of a cell with identified local intensity maxima marked in white. Scale bar B and C 10 μm, scale bar B’ and C’ 1 μm (C and C’ are adjusted for expansion), scale bar D 2 μm.

We imaged both fixed, unexpanded cells (Figure 3B) and expanded cells (Figure 3C) using an Airyscan microscope using a 40 × 1.2 NA water immersion objective. In the expanded samples, the non-homogenous distribution of NKA in the apical membrane was more clearly resolved than in the unexpanded samples.

To quantitatively describe the NKA organization, we analyzed the labeled protein distribution in our ExM images (Figure 3D). This analysis identified distinct aggregates in the expanded gel with sizes ranging from 150 to 600 nm, with a mean diameter of 286 nm. The lower bound of this range (150 nm) is constrained by the diffraction limit of our imaging setup. After correcting for the measured 4.3-fold expansion factor, these values correspond to a physical size range of ~35–140 nm. Taken into consideration that this number is still diffraction limited, it is in good agreement with previous super-resolution studies using SMLM and STED, which reported NKA cluster sizes in the 20–50 nm range (7,15,16).

We next tested the possibility of performing two-color labeling by combining a pre-expansion labeling step with a post-expansion one. It is possible to perform a first CuAAC labeling reaction with AF488-picolyl azide before fixation and a second reaction with AF647-picolyl azide after the sample has been denatured in the gel. Because the expansion protocol permeabilizes the cell, the pre-expansion AF488 labeling is limited to the plasma membrane, whereas the post-expansion AF647 labeling targets both extracellular and intracellular ncAAs (Supplementary Figure 1).

To validate the specificity of this intracellular, post-expansion labeling, we performed a control experiment in which NKA β_1_ WT or NKA β_1_ L64TAA was transfected together with the tRNA/aaRS pair for ochre suppression. Gels were labeled after denaturation with AF488-picolyl azide. Both cell types exhibited similar levels of intracellular fluorescence after post-expansion labeling, indicating that this signal was non-specific (Supplementary Figure 2). Furthermore, we tested if the SPIEDAC labeling chemistry could be performed after denaturation but found that its reactive handle (TCO*K) was no longer reactive after the gelation and denaturation process (not shown).

### Two color biorthogonal labeling of NKA subunits α_1_ and β_1_ in expanded cells

SPIEDAC and CuAAC click-chemistries can be combined for distinct two-color fluorescent labeling of ProK and TCO*K site-specifically incorporated into cell membrane proteins(17). For this we combined ochre suppression by M15_UUA_/*Mma*PylRS for ProK incorporation with amber suppression by hyb*_CUA_/*G1*RS YA for axial *trans*-cyclooct-2-ene-L-lysine (TCO*K) incorporation. We have successfully used this approach previously to label NKA subunits α_1_ and β_1_ with two fluorophores for quantification of NKA by FRET/FCS (3).

To perform two color expansion microscopy, we co-transfected HEK293T cells with NKA α_1_ T121TAG and NKA β_1_ L64TAA with both tRNA/aaRS pairs. AF488-picolyl azide was combined with Abberior STAR 635-tetrazine to spectrally separate the fluorescence emission. Both dyes were preserved throughout the expansion process and membrane expression of the two fluorescently labeled subunits could be detected (Figure 4A and A’). Unlike Abberior STAR 635-tetrazine, AF647-tetrazine was quenched during the expansion process (data not shown). Confocal Airyscan images of the lateral membrane revealed the distribution of α_1_ and β_1_ subunits with high contrast. We also measured the intensity of labeled α_1_ and β_1_ subunits along the highlighted lateral membrane region (Figure 4C), evaluated as normalized intensity ratios (I − I_min_) / (I_max_ − I_min_) and plotted a line profile (Figure 4 D). This visualization of the heterodimeric NKA is, to our knowledge, the first example of two-color super resolution expansion microscopy with GCE.

**Figure 4.**
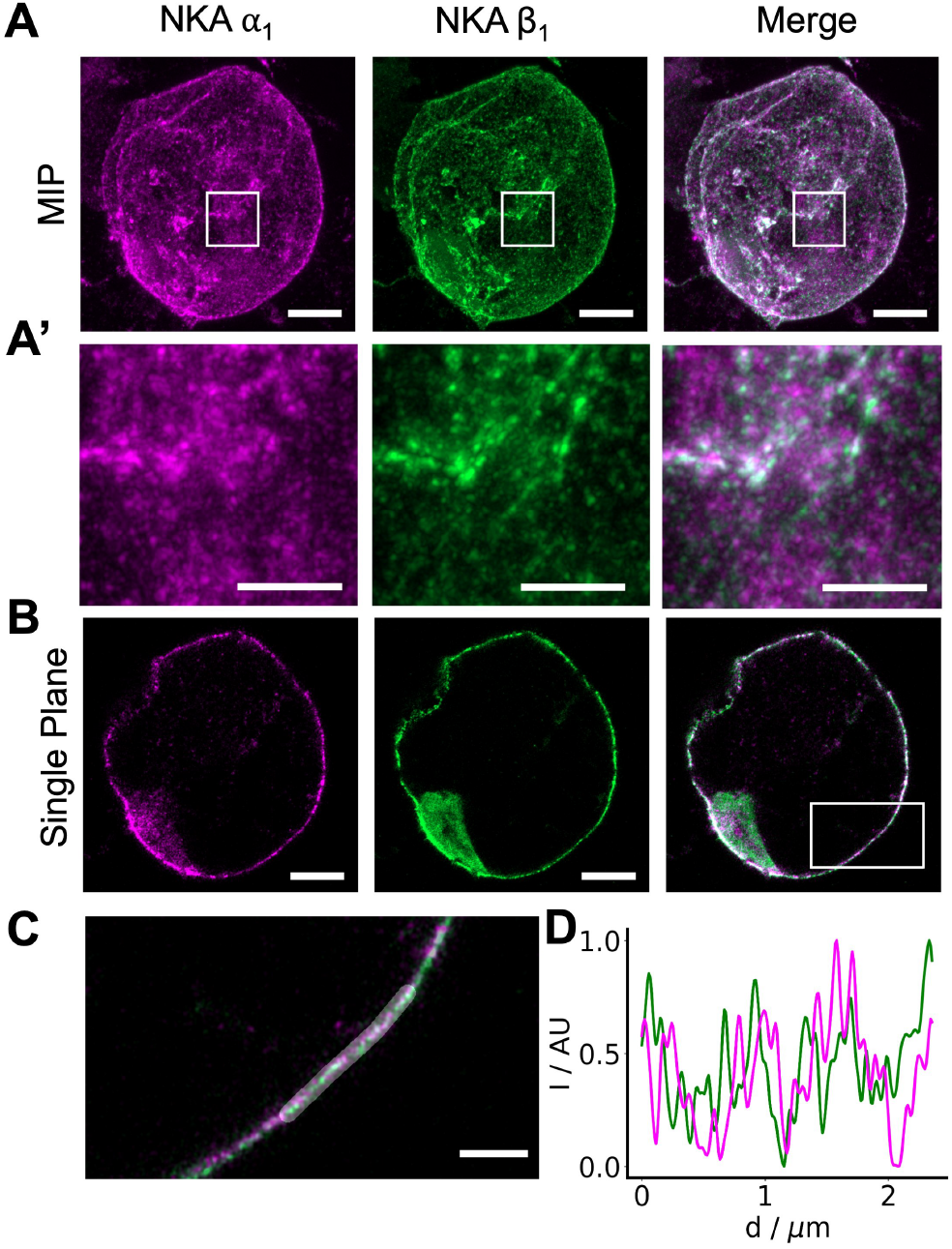
Two color biorthogonal labeling of NKA subunits α 1 and β 1 in expanded cells. Confocal z-stack (MIP) of an expanded HEK293T cell expressing NKA α1 T121TCO*K labeled with Abberior STAR 635-tetrazine (magenta) and β1 L64ProK labeled with AF488-picolylazide (green) (A). Boxed region from (A) shown in (A’). Single confocal plane through the center of an expanded HEK293T cell expressing NKA α1 T121TCO*K labeled with Abberior STAR 635-tetrazine (magenta) and β1 L64ProK labeled with AF488-tetrazine (green) (B). Boxed region from (B) shown in (C). Normalized line profile of highlighted membrane region in (C), showing heterogenous distribution of NKA α1 (magenta) and β1 (green) (D). Scale bars A and B 3 μm, scale bars A’ and C 1 μm (all adjusted for expansion).

## Discussion

In this study, we developed and applied a workflow combining GCE for site-specific labeling with ExM for super-resolution imaging. This approach enabled us to perform dual-color visualization of the NKA α_1_ and β_1_ subunits in HEK293T cells, providing nanoscale information on the enzyme’s distribution using a conventional confocal microscope. The primary motivation for this work is the fundamental challenge of determining the organization of proteins within the cell membrane. Many membrane proteins, including NKA, are proposed to exist not just as monomers, but also as dimers, higher-order oligomers, or organized in larger clusters (2–6). Distinguishing between these states is critical for understanding their function, but the nanometer-scale distances involved are far below the diffraction limit of conventional light microscopy. Therefore, resolving the true organization of such proteins demands super-resolution techniques that can achieve the highest possible localization precision.

A central advantage of GCE-based labeling is the minimization of “linkage error.” Common methods for fluorescence labeling using antibodies can add up to 20 nm of uncertainty to a protein’s position, a distance that can make it impossible to discern between a true dimer and two nearby monomers. Our approach reduces this to below 1 nm. Beyond labeling, the chemical fixation process itself poses a risk. Fixatives like paraformaldehyde (PFA) can induce artificial cross-linking of membrane proteins, potentially creating clusters that are not present in live cells (19–22). This possibility of fixation-induced artifacts must be considered when interpreting any super-resolution data on protein organization.

Stop codon suppression for fluorescent labeling is attractive, but also presents several challenges: Suppression of endogenous amber and ochre codons leads to read-through and ncAA insertion into unintended proteins, while inefficient suppression might result in truncated target proteins (23). In our case, the stop codon’s proximity to the mRNA’s 5’ end make it unlikely that truncated NKA subunits would fold correctly and be transported to the plasma membrane.

To avoid background from off-target ncAA incorporation, we used a membrane-impermeable dye to label extracellular domains. The validity of this strategy was confirmed with a control experiment. When we performed the click reaction after cell denaturation and permeabilization, we observed a bright, non-specific intracellular staining that was present in both wild-type and ncAA-expressing cells. This confirms the presence of an intracellular off-target signal, possibly from nuclear labeling of charged tRNAs (24), and validates that our extracellular-only strategy was essential for obtaining a clean signal.

An important step for the quality of labeling was to ensure that the fluorescent labels could survive the chemical treatments of the ExM protocol. We found that the cyanine-based dye AF647-tetrazine was quenched by free radicals during gel polymerization. In contrast, AF488-tetrazine and the rhodamine-based dye Abberior 635-tetrazine were well-preserved throughout the expansion process. This underscores that fluorophore stability must be empirically determined. To overcome quenching, it may be possible in the future to chemically modify sensitive dyes to protect them (25).

The aqueous hydrogel has a refractive index (RI) close to that of water (~1.33). Using a standard oil immersion objective (RI ~1.51) creates an RI mismatch that induces spherical aberrations, degrading the image (26). It is therefore important to use a water immersion objective to match the sample’s RI, preserving image quality and enabling accurate 3D data acquisition.

While STED and ExM offer impressive resolution, it remains insufficient to definitively resolve individual NKA subunits within the plasma membrane, especially if the protein is highly clustered. Our expansion microscopy data revealed incomplete colocalization of the α_1_ and β_1_ subunits. Since both subunits are required for transport to the plasma membrane, we hypothesize that a proportion of the transfected, labeled subunits formed functional complexes with the pool of endogenous, unlabeled NKA subunits. This unlabeled pool needs to be considered when measuring protein cluster size from super-resolution data.

In conclusion, our work establishes a robust framework for multiplexed super-resolution imaging that addresses key challenges of labeling precision and fluorophore compatibility. Although limitations (such as the efficiency of ncAA incorporation and potential for minor expansion-induced distortions) exist, this method can be broadly applied to investigate the nanoscale organization of a wide variety of protein complexes using widely accessible confocal microscopes. Future refinements in expansion chemistry could further improve resolution, bringing Ångström-scale structural biology within the reach of fluorescence microscopy.

## Acknowledgements

We thank Abberior GmbH for kindly providing Abberior STAR 635 and Abberior STAR RED tetrazine dyes. We also thank Linnea Nordahl, Katja Schach and Philipp Graef for laboratory assistance, Jonatan Alvelid for image analysis, and Evgeny Akkuratov and Johannes Heimgärtner for their contribution in the early stages of this study.

---

**Figure S1.**
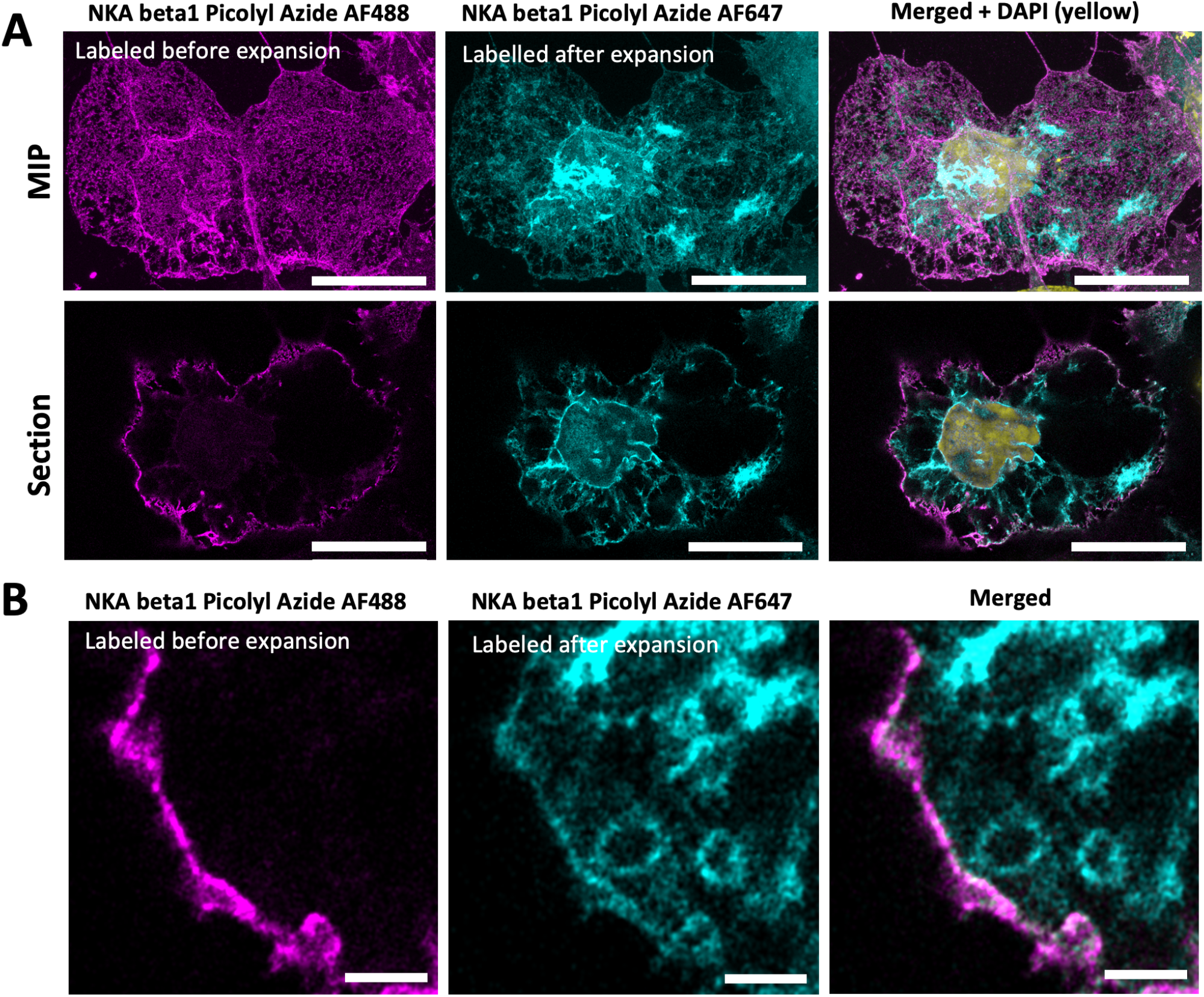
CuAAC labeling can be performed after denaturation to label intracellular ncAAs. Confocal image of an expanded HEK293T cell expressing NKA β_1_ L64ProK labeled before fixation with AF488-picolyl azide (magenta) and after denaturation with AF647-picolyl azide (cyan), top row: MIP, bottom row: sing section **(A)**. Zoomed confocal image of an expanded HEK293T cell expressing NKA β_1_ L64ProK labeled before fixation with AF488-picolyl azide (magenta) and after denaturation with AF647-picolyl azide (cyan) (**B**). Scale bar (A): 10 μm, scale bar (B) 4 μm (adjusted for expansion).

**Figure S2.**
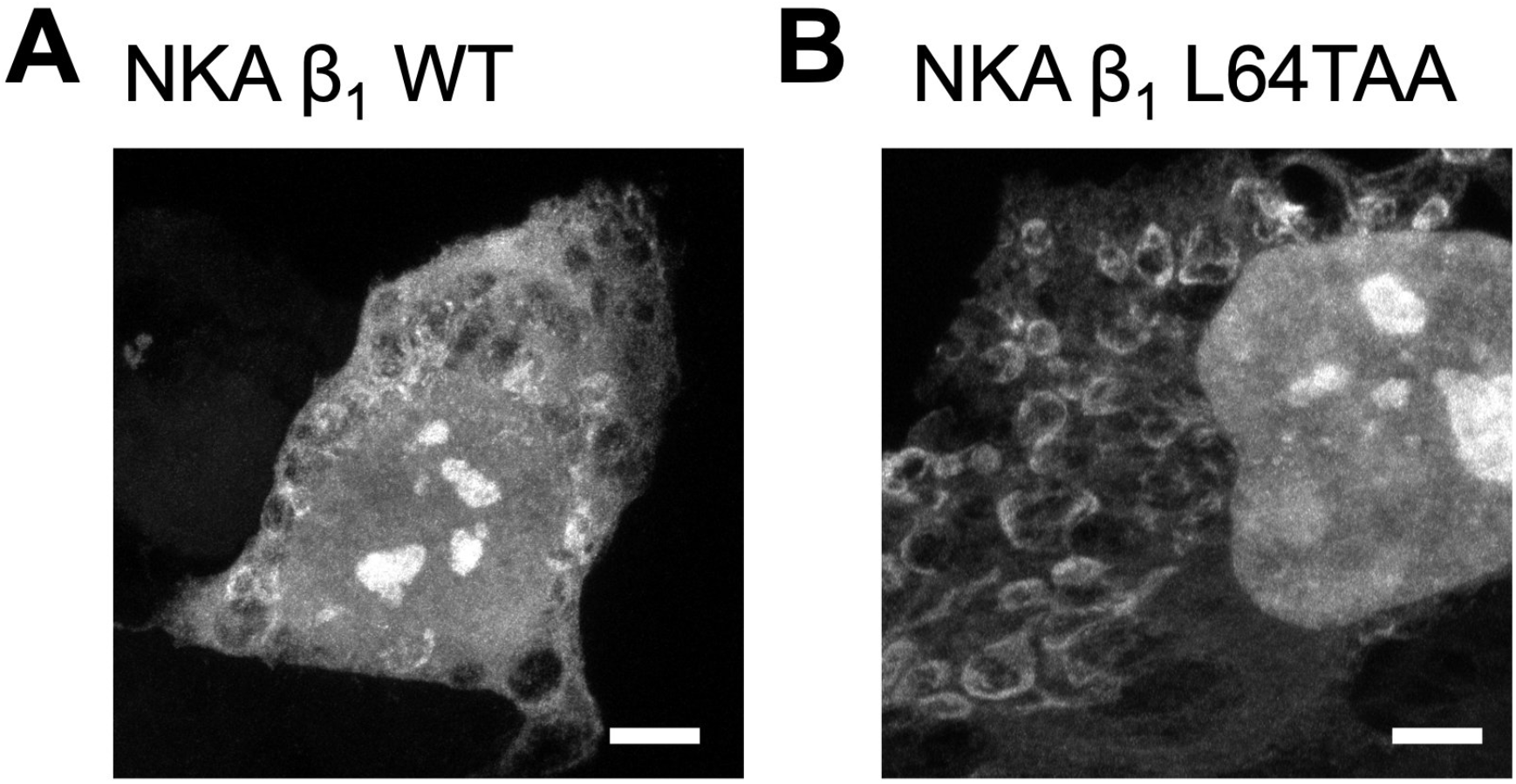
Post-denaturation labeling of both NKA β 1 WT and β 1 L64TAA transfected cells shows intracellular structures. Confocal stack (MIP) of HEK293T cells transfected with M15_UUA_/*Mma*PylRS and NKA β_1_ WT, grown with ProK, expanded, and CuAAC labeled with AF488-picolyl azide after denaturation (A). Confocal stack (MIP) of HEK293T cells transfected with M15_UUA_/*Mma*PylRS and NKA β_1_ L64TAA, grown with ProK, expanded, and CuAAC labeled with AF488-picolyl azide after denaturation (B). Scale bar 10 μm.

## Notes

### Competing Interest Statement

The authors have declared no competing interest.

### Funding

- Swedish Research Council: 2020-05347, 2023-00163

